# Fully-automated and ultra-fast cell-type identification using specific marker combinations from single-cell transcriptomic data

**DOI:** 10.1101/812131

**Authors:** Aleksandr Ianevski, Anil K Giri, Tero Aittokallio

## Abstract

Single-cell transcriptomics enables systematic charting of cellular composition of complex tissues. Identification of cell populations often relies on unsupervised clustering of cells based on the similarity of their scRNA-seq profiles, followed by manual annotation of cell clusters using established marker genes. However, manual selection of marker genes is a time-consuming process that may lead to sub-optimal annotation results as the selected markers must be informative of both the individual cell clusters and various cell types present in the complex samples. Here, we developed a computational platform, termed ScType, which enables data-driven, fully-automated and ultra-fast cell-type identification based solely on given scRNA-seq data, combined with our comprehensive cell marker database as background information. Using a compendium of six scRNA-seq datasets from various human and mouse tissues, we show how ScType provides an unbiased and accurate cell-type annotation by guaranteeing the specificity of positive and negative marker genes both across cell clusters and cell types. We also demonstrate how ScType enables distinguishing between healthy and malignant cell populations, based on single-cell calling of single-nucleotide variants, making it a versatile tool for exploration and use of single-cell transcriptomic data for anticancer applications. The widely-applicable method is deployed both as an interactive web-tool (https://sctype.app), and as an open-source R-package, connected with a comprehensive ScType database of specific markers.

## Introduction

Accurate identification of distinct cell types in complex tissue samples is a critical prerequisite for elucidating the roles of cell populations in various biological processes including hematopoiesis, embryonic and intestinal development^1,2,3,4^. Traditionally, cell sorting and microscopic techniques have been extensively used to isolate cell types, followed by molecular profiling of the sorted cells using, for instance, mRNA or protein measurements^5,6,7^. Decades of research has led to several collections of cell-specific features, including expression of marker genes in various tissues, that are being used to distinguish various cell types in new tissue samples^8,9^. However, the entire process is manually tedious and technically challenging. Recently, single-cell RNA sequencing (scRNA-seq) has been established as a high-throughput approach to routinely chart diverse cell populations in tissue samples and to study various biological processes in disease and development^2,10,11,12^. The scRNA-seq technology has provided an unprecedented view of various cell types and it has become the leading technology in large-scale cell mapping projects such as the Human Cell Atlas^13^.

Identification of cell populations often relies on unsupervised clustering of cells based on their transcriptomic profiles, followed by cluster annotation using marker genes that are differentially-expressed between the clusters^14,15^. These marker genes are then manually inspected using available information in the literature or cell marker databases^8,9^ to assign cell-type labels to each detected cluster. However, the manual selection of cluster-specific marker genes is a time-consuming and error-prone task, since the marker genes are often (*i*) expressed in multiple cell clusters, and (*ii*) correspond to multiple cell types. In addition, the expression of *negative marker genes*, which provide evidence against a cell being of a particular type, should also be incorporated into the cell-type identification process. The cell annotation procedure is further complicated by the lack of curated cell marker databases that include both known and *de novo* positive and negative markers to annotate cell-types with confidence. For example, selection of CD44 as marker gene may compromise the accuracy of cell annotation as CD44 is expressed in various immune cell populations.^8^ Another popular approach for cell-type assignment is to utilize a reference dataset, a collection of previously annotated cell types in single-cell data, to train a classification algorithm and to apply it to new single-cell datasets. However, such supervised approaches require that the reference and new datasets resemble each other, which often pose a problem in scRNA-seq studies^16^.

One important application of single-cell characterization is to design personalized treatments that selectively target malignant cell types in a patient-derived sample, while avoiding severe inhibition and toxic effects on healthy cells^17^. In cancers and other complex diseases, monotherapy resistance often emerges and requires multi-drug co-inhibition of various disease-and resistance-driving cell populations. We recently demonstrated how our comprehensive ScType marker database helped an AI-guided identification of personalized drug combination therapies for patients with refractory acute myeloid leukemia (AML), which led to synergistic co-inhibition of leukemic cell subpopulations that emerged in various stages of the disease pathogenesis and treatment regimens^18^. These cancer-selective and patient-specific combinations were shown to be relatively less toxic to lymphoid cells (non-malignant cells in the AML case), thereby increasing their likelihood for clinical translation. However, how to accurately distinguish between multiple malignant and non-malignant cell populations for targeted treatments remains a translational challenge and requires both systematic and highly selective strategies that are applicable to various diseases and tissue types. In many applications, reference single-cell data and cell-type annotations are not available, rather the cell population identification needs to be done individually for each patient sample.

To solve these challenges, we developed a computational ScType platform (marker database and cell-type identification algorithm), which requires only a single scRNA-seq dataset for accurate and unsupervised cell-type annotation (Fig. 1a). The unbiased yet selective cell-type annotation is achieved by compiling the largest database of established cell-specific markers (ScType database), and by ensuring the specificity of marker genes across both the cell clusters and cell-types (ScType algorithm, see Fig. 1b-c). We carried out a systematic benchmarking of ScType and related methods across 6 scRNA-seq datasets from 4 human and 2 mouse tissues, and showed that ScType platform correctly annotated a total of 72 out of 73 cell-types (98.6% accuracy), including 8 newly-reannotated cell-types that were incorrectly or non-specifically annotated in the original studies. In addition, ScType implements a single-cell single-nucleotide variant (SNV) calling option to distinguish between malignant and non-malignant cells (Fig. 1a), demonstrated here using scRNA-seq data from AML patient samples. The ScType platform is implemented as an open-source and interactive web-tool (https://sctype.app), connected to the ScType marker database, to enable ultra-fast and fully-automated cell-type annotation.

**Figure 1.**
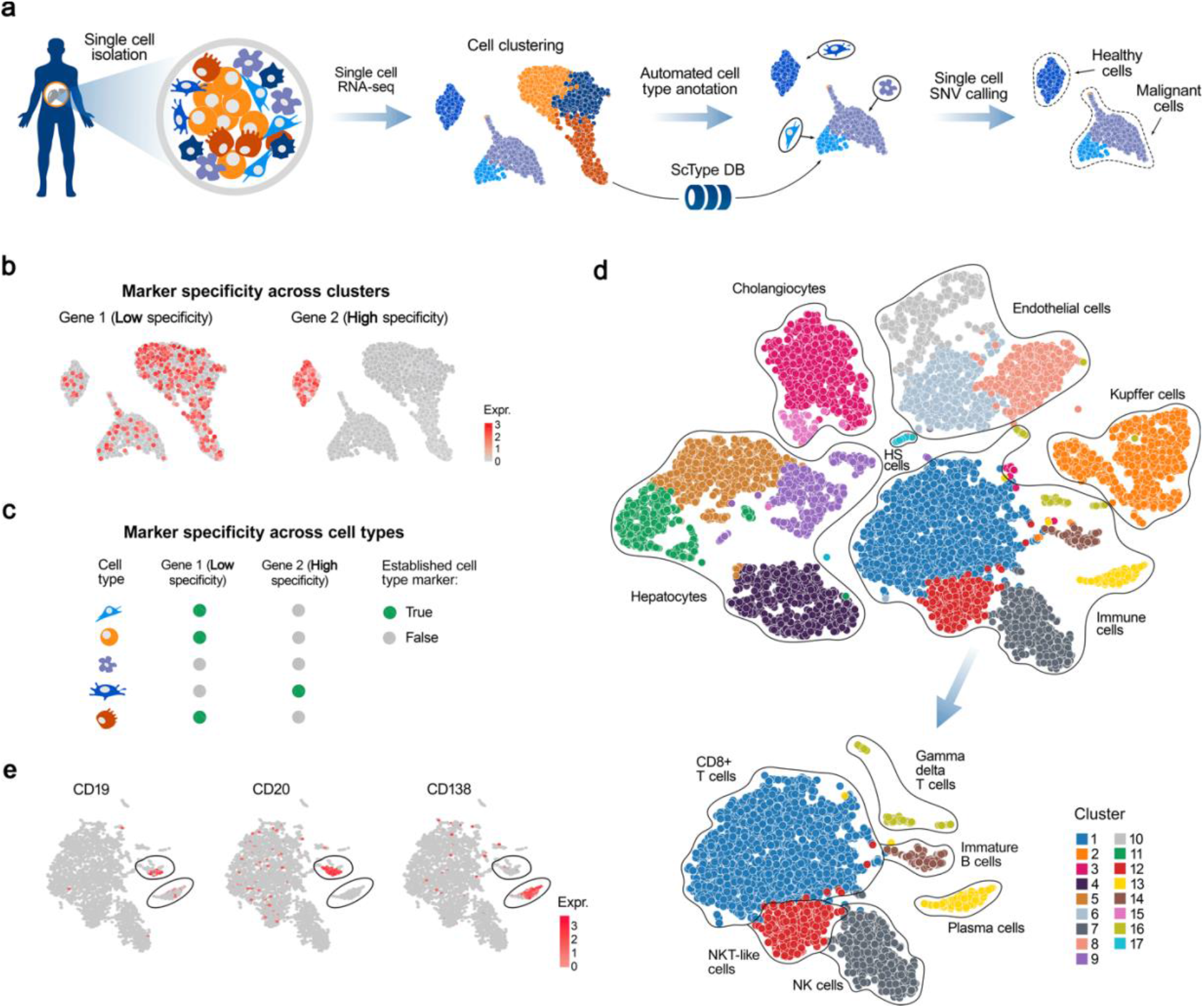
A schematic view of cell-type annotation using ScType. **(a)** ScType requires only the raw or pre-processed single-cell transcriptomics dataset(s) as input. ScType implements options for additional quality control and normalization steps, where needed, followed by unsupervised clustering of cells based on scRNA-seq profiles. The results here are based on the Louvain clustering; however, also SC3, DBSCAN, GiniClust and k-means clustering options are available in ScType (see Methods). In the next step, ScType performs a fully-automated cell-type annotation using an in-built comprehensive marker database. Finally, ScType implements novel options for somatic single-cell SNV calling to distinguish between healthy and malignant cell populations. **(b-c)** ScType algorithm guarantees that the marker genes show specificity both across clusters and cell types for accurate unsupervised cell-type annotation with high cell subpopulation selectivity. **(d)** UMAP example of automated cell subtype identification by ScType in the liver atlas dataset, where it automatically labelled the same cell-types as assigned manually in the original study.^10^ **(e)** Based on the information that plasma cells do not express common B-cell markers, such as CD19 and CD20, but instead express CD138, ScType enhanced the resolution of cell-type annotations of two cell clusters, which were jointly annotated as B-cells in the original study, by segregating them into immature B-cell and plasma (B) cell types (lower UMAP plot of panel **d**).

## Results

### ScType improves annotation of cell-types using solely scRNA-seq data

We first investigated the performance of ScType by re-analyzing a published scRNA-seq study of human liver cells^10^. Using only the raw scRNA-seq data from the liver atlas dataset, ScType identified 17 clusters and correctly assigned them to 11 identified liver-related cell types that were manually annotated in the original study (Fig 1d). This demonstrates the benefits of the comprehensive marker databases and the accuracy of the fully-automated annotation process. Additionally, ScType was able to automatically distinguish between two closely-related cell populations of B-cells (immature and plasma B cells) that were not differentiated in the original manuscript^10^. This segregation between immature and plasma B cells was done based on the information in the ScType database that plasma cells do not express common B-cell markers, such as CD19 and CD20, but instead they express CD138 (Fig. 1e)^19^.

Next, we re-analyzed another published scRNA-seq data of mouse retinal cells (Fig. S1a)^20^. ScType automatically identified three closely-related cell populations of amacrine cell types (GABAergic, glycinergic and startbust), which were originally-identified by an extensive and deep analysis of selectively-expressed markers^20^. Furthermore, ScType correctly distinguished between the two subtypes of bipolar cells – rod (expressing PRKCA^21^ and CAR8^22^, Fig. S1b) and cone (expressing SCGN^23^, Fig. S1b) bipolar cells, which were manually assigned to a single group in the original study, therefore enhancing the resolution of the cell-type annotation. Taken together, these results indicate that ScType enables a fully-automated prioritization of highly-specific markers for accurate annotation of even rare cell-types with distinct and selective molecular features.

### Systematic evaluation of ScType across human and mouse scRNA-seq datasets

To investigate the wider applicability of the automated method, we next benchmarked the performance of ScType in terms of its ability to automatically assign cell-types in comparison to the cell-type annotations given by the original authors of additional four (six in total) published scRNA-seq studies. We further utilized all the six datasets to compare ScType performance against other recent cell-type annotation methods in terms of their accuracy and running time. The RNA-seq datasets used in the benchmarking originated from various tissues, including human liver^10^, pancreas^24^, peripheral blood mononuclear cells (PBMCs)^25^, brain^26^, as well as mouse lung^27^ and retina samples^20^. These scRNA-seq datasets were utilized to investigate the performance of ScType and the related methods in the context of various sequencing platforms, tissues types and organisms.

Among the six scRNA-seq datasets from various human and mouse tissues, ScType correctly annotated a total of 72 cell types out of 73 cell-types (98.6% accuracy), including 8 correctly reannotated cell-types that were originally incorrectly or non-specifically annotated (Fig. 2a). The only cell-type ScType was unable to automatically label as known was fetal cells in the human brain dataset, as there are no fetal cell markers available for human brain in the current version of the ScType database. However, ScType correctly identified all the other cell populations of the human brain tissues (oligodendrocytes, astrocytes, microglial cells, neurons, endothelial and oligodendrocyte precursor cells), according to the annotations made in the original study^26^. Furthermore, ScType was able to refine the originally-annotated neuron cell population into cholinergic (expressing SLC17A7)^28^ and glutamatergic (expressing ACHE)^29^ subtypes.

**Figure 2.**
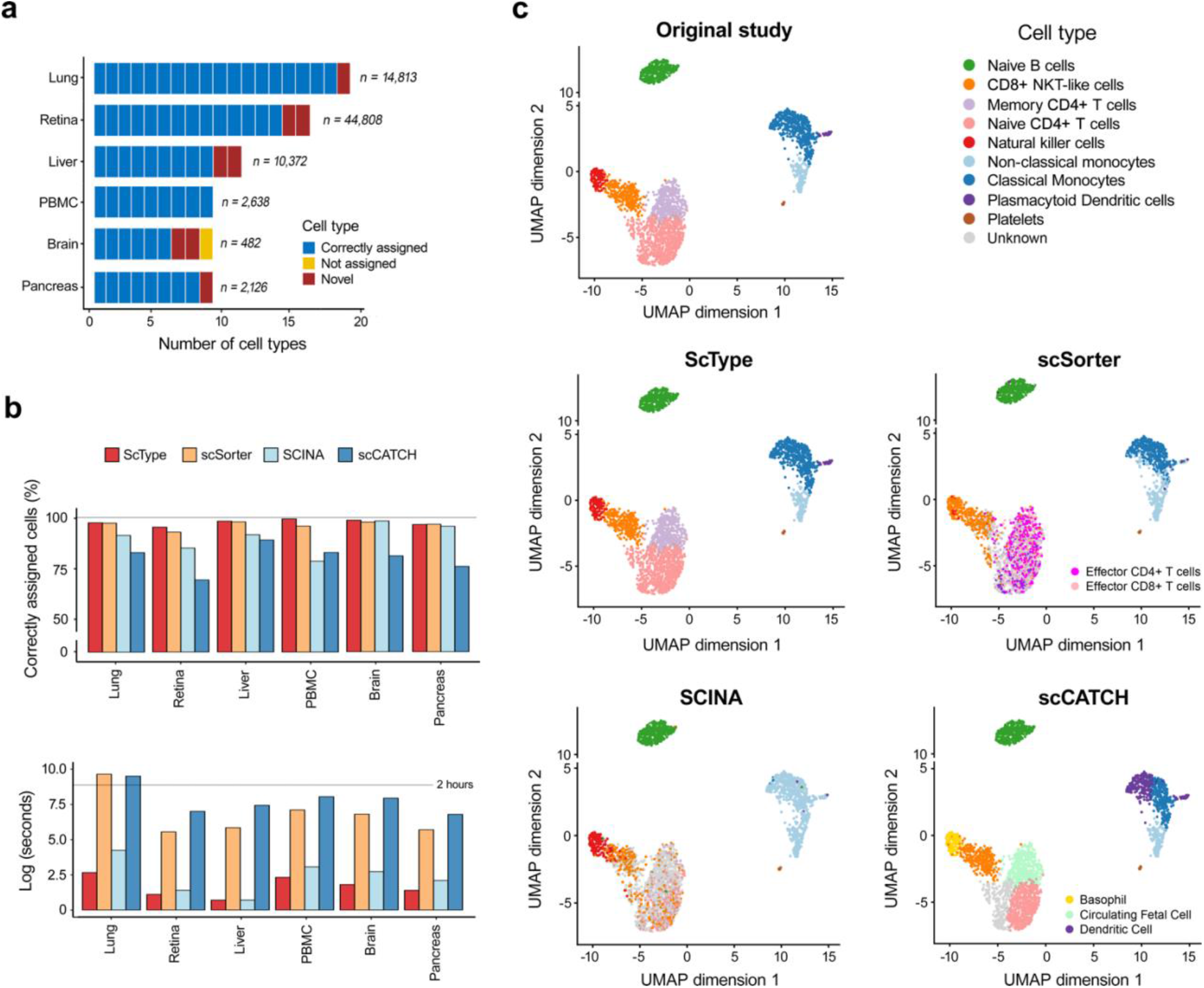
ScType performs ultra-fast and accurate cell-type annotation across various tissues. **(a)** The overall performance of ScType across six human and mouse scRNA-seq datasets. ScType automatically assigned cell-types according to the original studies, and it also correctly reannotated five cell types in the human brain, liver and pancreas tissues, compared to the original studies (marked as novel cell types). ScType labelled only single cluster (fetal cells) as unknown cell-type in the human brain dataset (marked as not assigned). Similarly, in the mouse lung and retina datasets, ScType enabled automated identification of all the cell types, and it also correctly reassigned three novel mouse cell types. **(b**) Comparison of ScType with three recently developed cell-type annotation methods in terms of percentage of correctly annotated cells (upper panel) and running time (lower panel). **(c)** Detailed cell-type annotations of the human PBMC dataset by the methods under comparison.

Next, we compared ScType against the state-of-the-art cell-type annotation methods with reported (i) highest accuracy (scSorter^30^ that was recently shown to outperform several popular tools such as Garnett^31^ and CellAssign^32^), (ii) shortest running time (SCINA^33^), and (iii) full-automated process (scCATCH^34^). For an unbiased comparison, we provided scSorter and SCINA with our in-built database markers, while scCATCH utilized its own integrated marker database. Overall, ScType correctly annotated more than 94% of the cells in each dataset (Fig. 2b, upper panel), outperforming the other algorithms in 5 out 6 datasets. We note that the differences between ScType and the next best performing method scSorter were not large, as both methods showed a high accuracy in all the datasets, but importantly ScType was more than 30 times faster than scSorter (Fig. 2b, bottom panel). In particular, ScType showed almost perfect accuracy in the challenging human PBMC dataset (Fig. 2c), where there are multiple closely-related subtypes. In contrast, scSorter and scCATCH did not identify natural killer cell population, and they incorrectly identified several T cell subtypes, while SCINA was not able to distinguish between the two monocyte subpopulations as well as several subsets of T cells (Fig. 2c).

These benchmarking results indicate that ScType enables ultra-fast and highly accurate separation even between closely-related subtypes by utilizing the novel concepts of marker gene specificity across both cell clusters and cell types, along with negative marker genes (e.g., effector T cells are known to be CCR7 negative^35^).

### Single-cell SNV calling distinguishes between healthy and malignant cell types

To enable genetic analyses in cancer applications, we further implemented an option for single-nucleotide variation (SNV) calling directly from the scRNA-seq data. As an example, we re-analyzed the scRNA-seq transcriptomic profile and cell-type composition of an AML patient sample from our recent study^18^ using the ScType platform (Fig 3a). After performing automated single-cell SNV calling (see Methods), we investigated whether the number of SNVs within a cell-type could distinguish between healthy and cancerous populations present in the patient sample (Fig. 3b). More specifically, ScType quantified the percentage of cells in a cell-type above the median SNV in the cancer consensus genes across all cells within the sample (ScType_SNV score). As expected, we observed a higher SNV score in the CD34+ progenitor (HSC/MPP) cells and CD34+ interferon-stimulated gene (ISG)+ blast cells, as compared to CD24+ CD66b+ neutrophils and memory CD8+ T cells that are usually considered as non-malignant cell types in AML^11^ (Fig. 3c). As another validation for correctly distinguishing normal cells from malignant cells, we considered aneuploidy (unbalanced number of chromosomes), which is common for most human tumors^36^. To identify aneuploidy, we incorporated the recent Bayesian segmentation approach, CopyKAT^36^, that classified majority of CD24+ CD66b+ neutrophils and memory CD8+ T cells as diploid cells (Fig. 3d), suggesting their non-malignancy. These two validations demonstrated how ScType correctly assigned CD24+ CD66b+ neutrophils and memory CD8+ T cells as non-malignant cells (ScType SNV score < 20 and the majority of cell within the cell-type are classified as diploid cells, Fig 3e).

**Figure 3:**
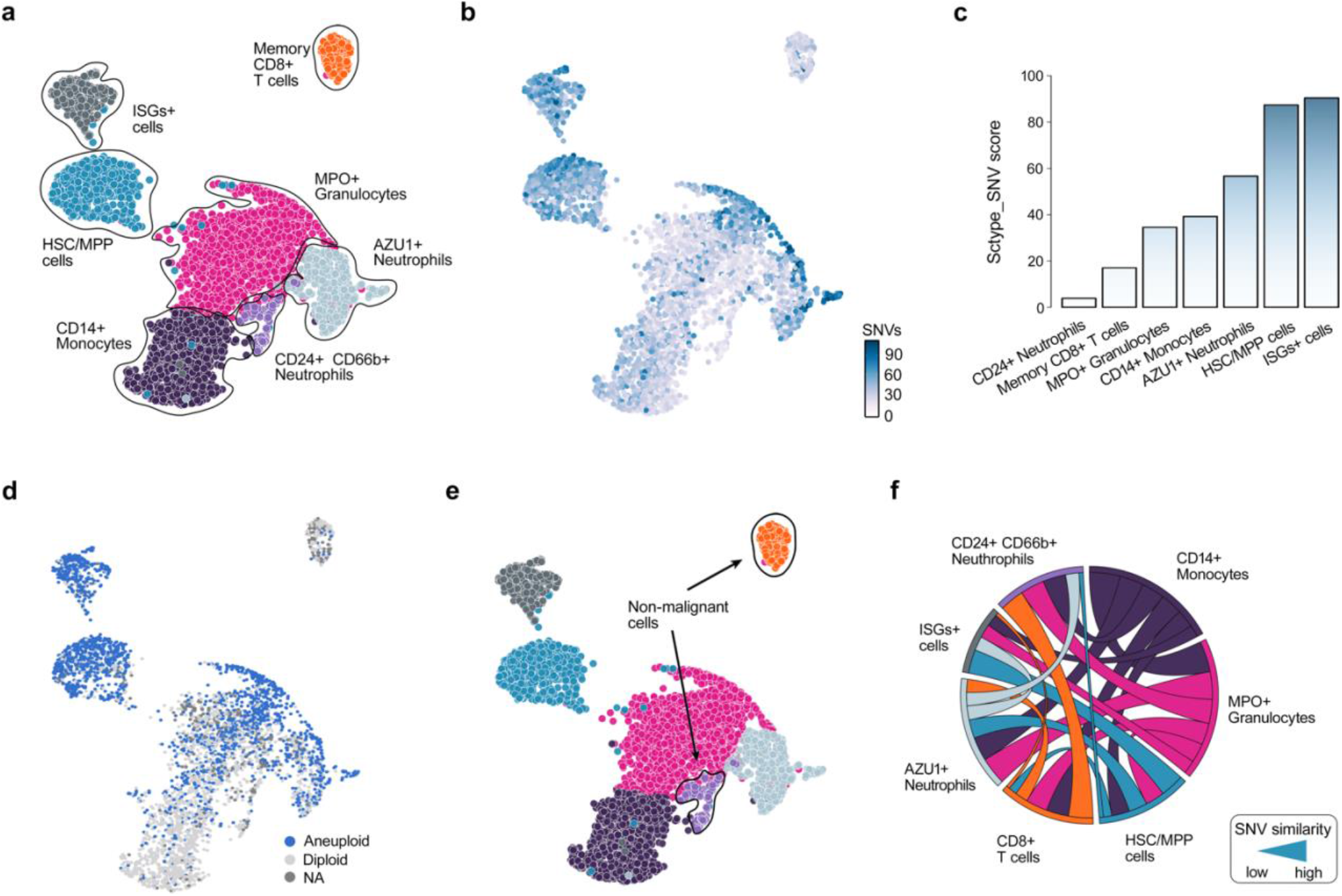
ScType enables SNV calling directly from the scRNA-seq profiles. **(a)** UMAP plot of the cell types in the AML patient sample automatically annotated by ScType. **(b)** Distribution of somatic SNVs in the cancer consensus genes across the various cell types of the patient. **(c)** ScType SNV score summarizes the number of point mutations in the cancer genes for each cell type, shown as the percentage of SNVs above the median SNV value within the particular cell cluster. **(d)** UMAP showing aneuploid and diploid cell classification based on Bayesian segmentation approach CopyKAT^36^. **(e)** ScType assigns cell types as non-malignant when the ScType_SNV score is below 20 and more than 50% of cells within the cell-type are classified as diploid. **(f)** Chord diagram shows the associations between different cell types in terms of the similarity of the SNVs in the cancer genes (i.e., occurrence of common SNVs, see Methods for details). The width of a connection corresponds to the degree of SNV similarity between cell-types, while the connection color indicates a specific cell-type as shown in UMAP plot in Figure 3a.

To further investigate the ScType cell population classification, we studied the associations between the various cell-types based on the occurrence of common SNVs in the cancer consensus genes, and observed that non-malignant cell-types (i.e. memory CD8+ T cells and CD24+ CD66b+ neutrophils) were closely similar to each other, while showing almost no SNV similarity with the malignant cell types (e.g. HSC/MPP and ISG+ blast cells, see Fig. 3f). These results demonstrate how the ScType platform enables one to distinguish between malignant and non-malignant cell populations, based directly on scRNA data from a given patient sample, which is critical for the selection of safe and effective targeted treatment regimens for individual AML patients. For other cancer types and malignancies, the platform similarly supports automated options for the marker selection and cell population classification into healthy and diseased cells. In addition, ScType enables the visualization of the genome-wide copy number profiles from scRNA-seq data using CopyKAT to identify larger-scale copy number alterations (CNAs), such as somatic gains or deletions of large segments of chromosomes (see Fig. S2 as an example in the AML patient sample). In the downstream analyses, the identified CNAs may explain difference in the cellular phenotypes of specific cell types and subclones, including their apoptotic potential or drug sensitivity.

## Discussion

We presented ScType, a fully-automated platform for cell-type identification that enables accurate and ultra-fast single-cell-type annotations based solely on the given scRNA-seq data, using our comprehensive ScType marker databases as background information. To the best of our knowledge, ScType is currently the only unsupervised method that makes use of the marker gene specificity across both cell clusters and cell types to automatically identify highly-specific positive marker genes, along with negative marker genes to provide evidence against cells being of a particular cell-type for cell-type selective annotations. To promote its wide application, either as a stand-alone tool or together with other popular single-cell data analysis software (e.g., Seurat^37^, MAST^38^, PAGODA^39^), we have deployed ScType both as an interactive web-platform (http://sctype.app), and as an open-source R implementation (http://sctype.app/source_code.php).

We anticipate the ScType platform will accelerate unbiased phenotypic profiling of cells when applied either to large-scale single-cell sequencing projects or smaller-scale profiling of patient-derived samples. For example, the integrative marker information in the ScType database may enable the identification of rare cell subtypes that have distinct combinations of molecular markers, suggesting specific functions and phenotypic profiles. We recently demonstrated how the ScType database provided information for patient-tailored identification of cancer-selective combinatorial therapies for relapsed AML patients, each with different genetic background and resistance mechanisms^18^. The ScType annotation algorithm, together with the novel methods to distinguish between malignant and non-malignant cells, are expected to enable design of targeted treatment regimens also for other cancer types.

The existing computational methods for automatic identification of cell types can be broadly categorized into two groups: (1) supervised methods that require carefully-annotated training datasets labelled with correct cell populations to train the classifiers (e.g. CaSTLe^40^ and ACTINN^41^ that annotate cell types based on pre-defined reference set of cells without the need of cell marker input), and (2) a prior knowledge-based methods that require either a marker gene set or a pre-trained classifier for the selected cell populations (e.g. scSorter^30^, SCINA^33^, and scCATCH^34^). The supervised methods may have severe limitations when annotating especially rare populations of cells, due to lack of reference data to train the machine learning algorithms. Furthermore, supervised methods are notoriously time-consuming to train, as well as error prone to technical artifacts in the training data, which affect their prediction ability for new scRNA-seq data^42^.

Similarly, the prior knowledge-based cell classification approaches have certain limitations. For instance, their performance heavily depends on the available gene lists provided as markers for each cell type, typically obtained from manual literature search or matching to marker databases that are still suboptimal both in coverage and specificity. Ideally, one would like to use an appropriate number of specific markers to achieve a maximally accurate cell-type classification. However, most existing methods utilize a limited number of markers, thereby potentially masking the identification of a subpopulation of cells that do not express the selected marker genes. Furthermore, the use of inconsistent cell-type markers across experiments and laboratories may compromise the reproducibility of the findings^42^. These caveats become even more pronounced when the number of cell types and samples increases, thus preventing fast and reproducible annotations. It has been therefore argued that prior information does improve the automated cell-type identifications^32,42.^

ScType implements a number of improvements compared to the existing cell-annotation tools. Our unsupervised approach outperformed the prior knowledge-based methods scSorter^30^, SCINA^33^, and scCATCH^34^, which were recently shown to enable accurate annotation of multiple cell types^42^. Another group of supervised methods, such as CaSTLe^40^, ACTINN^41^, SingleR^43^ and CHETAH^44^, utilize reference bulk or single-cell transcriptomic data for cell-type predictions, and therefore require comprehensive, manually-annotated and high-quality reference datasets for training; furthermore, these methods do not allow identification of novel cell-type marker genes. In contrast, ScType requires neither reference scRNA-seq datasets nor manual selection of marker genes; instead, all the background information for established or *de novo* markers comes from the ScType database that is to date the most comprehensive database of specific markers for human and mouse cells.

In comparison with many other computational approaches that require manual interference,^31,32^ ScType takes a fully data-driven approach, and it annotates the cell-types at once in a totally unsupervised manner. The only input needed for the ScType tool is the raw sequencing data file, although uploading of pre-processed scRNA-seq data is also an option. This saves considerable time and costs in the scRNA-seq analysis, especially when searching for cell-types in a tissue that involves a large variety of cell-types with similar transcriptomic profiles (e.g. bone marrow samples from mixed lineage leukemia subjects). The running time of ScType is also orders of magnitude faster than the supervised methods. Furthermore, ScType implements options for SNV and CNV calling from the raw scRNA-seq profiles of individual samples. The users may compare SNV levels across different cell types, and study associations between cell clusters based on their SNV load.

Using six scRNA-seq datasets from the human and mouse tissues, we demonstrated that ScType provides scalable and accurate identification of cell-clusters, and it is compatible with data formats from various sequencing techniques (e.g. Drop-seq and Smart-seq). These benchmarking results against the existing cell annotation approaches indicated that ScType is widely-applicable to various biomedical problems, and it provides fast and accurate cell-type classifications. Furthermore, we expect that the comprehensive ScType database will lead to the development of new and improved cell-type detection methods, as well as accelerate the implementation of single-cell pipelines for translational applications, such as monitoring of therapy resistant cancer cell sub-populations and designing of targeted combinatorial therapies to overcome the monotherapy resistance in cancer patients, which require fast and automated analyses for real-time clinical implementation.

In conclusion, ScType provides automated cell-type identification using its own in-built database, as well as identification of malignant cell populations and cancer targets based on SNV calling and aneuploidy identification, requiring only raw scRNA-seq data as an input. As increased number of scRNA-seq datasets from various tissue types become available from the Human Cell Atlas and other projects, the accuracy and coverage of the ScType platform is expected to increase accordingly.

## Authors contributions

AI, AKG and TA conceived and planned the study. AI developed the method, implemented the ScType web-tool and collected and analyzed the data. AI compiled the ScType database with the help of AKG. AI prepared the figures for manuscript with the help of TA and AKG. TA and AKG supervised the study. All the authors wrote the manuscript and approved its final version.

## Acknowledgements

The authors thank Dr. Pirkko M Mattila, Dr. Jenni Lahtela and Bhiswa Ghimire for their valuable suggestions to improve the web-tool, and Olle Hansson for the cluster server machine to host the web-tool and the database. This work was supported by the Academy of Finland (grants 295504, 310507, 326238 and 344698 to TA), European Union’s Horizon 2020 Research and Innovation Programme (ERA PerMed JAKSTAT-TARGET), the Cancer Society of Finland (TA), the Sigrid Jusélius Foundation (TA), and the Norwegian Cancer Society (grant 216104 to TA).

## ONLINE METHODS

### ScType database construction

ScType database is the largest database to date of human and mouse cell-specific markers, compiled by integrating the information available in the CellMarker database (http://biocc.hrbmu.edu.cn/CellMarker/) and PanglaoDB (https://panglaodb.se), which are currently the two largest available databases for cell-type markers. In the CellMarker database, 13 605 cell markers for 467 cell types in 158 human tissues/sub-tissues and 9148 cell makers for 389 cell types in 81 mouse tissues/sub-tissues were manually collected and curated from more than 100 000 published papers^8^. In the PanglaoDB, 6631 gene markers mapping to 155 cell types have been identified by differential expression analysis in the particular cell types using single-cell data and a community-based crowdsourcing approach for curation of gene expression markers^9^. However, these two databases differ in the number of tissues, cell types and marker numbers, as well as in the way the markers have been assigned to each cell type. Therefore, we firstly converted the non-uniform gene IDs to approved gene symbols within and between the databases. Next, we removed the low evidence marker genes from the CellMarker database (i.e., genes having only one reference to support the cell-type marker), and genes that appeared in less than 5 clusters of specific cell-type from PanglaoDB. Additionally, we excluded genes showing no expression across all the datasets in PanglaoDB. Ultimately, we unified the cell and tissue naming from the two databases and excluded tissues comprising less than 5 cell types. Fifteen novel cell types with corresponding marker genes were added by manual curation of >10 papers to the current version of the compiled ScType database (https://sctype.app/database.php), as relatively few brain and eye tissue cell types were provided in the first version of the database. In total, the current version of the ScType database comprises 3980 cell markers for 194 cell types in 17 human tissues and 4212 cell markers for 194 cell types in 17 mouse tissues.

### Cell-type specificity of markers

Cell-type specificity (*S*) was calculated separately for each marker gene (*M*_*i*_) across the cell types, hence providing a quantitative measure of how frequently the marker maps to the cell-type uniquely within a particlar tissue (*t*) using the cell-type specificity score: 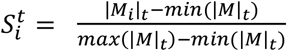, where *M* = (*M*_1_, …, *M*_*i*_, …, *M*_*m*_), and *m* is the total number of marker genes *M* present across all the cell types of the tissue *t*, and *i* is the index of each unique marker.

### ScType workflow options

ScType provides a complete pipeline for single-cell RNA-seq data analysis and cell-type annotation. We utilized Seurat v4.0^37^ for data processing and normalization. For clustering analysis, the default option is Louvain clustering based on a shared nearest neighbor graph (using FindClusters function with the resolution parameter set to 0.8 and 20 principal components given as input), which was used to generate the current results; however, also SC3^45^, DBSCAN^46^, GiniClust^47^ and k-means clustering options are available in ScType. The clusters are visualized using either principal components analysis (PCA), t-distributed Stochastic Neighbor Embedding (t-SNE), Uniform Manifold Approximation and Projection for Dimension Reduction (UMAP), Isomap^48^, Diffusion Map^49^, largeVis^50^ or by means of expression heatmaps. For the integrated multi-scRNA-seq dataset analysis, ScType uses FindIntegrationAnchors and IntegrateData functions from Seurat v4.0 that were shown to enable an effective identification of anchor correspondences across multiple single-cell datasets^37^. As a key unique component, ScType implements an ultra-fast and fully-automated cell-type identification, using highly-specific marker genes (see above), and allows the user to explore each gene’s contribution to cell-type annotations (see Fig. S1). Finally, Sctype implements options for SNV calling and aneuploidy identification, and it automatically suggests separation between non-malignant and malignant cell types (see below).

### ScType cell-type annotation

In order to assign each cell-type to a cluster (*p*), given the input scRNA-seq data (*X*) with *m* genes and *n* cells, ScType first standardizes each gene expression profile into z-score across all genes. Only positive and negative markers genes corresponding to different cell types of the specified tissues are considered (extracted either from ScType database or using user-provided custom cell-type gene sets as an alternative option). Then, each gene expression level is multiplied with its cell type-specificity 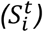 score: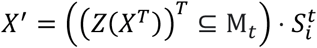, where *X*′ is a transformed scRNA-seq expression matrix of *n* cells and |M_*t*_| genes, M_*t*_ is the vector of marker genes of all cell types within the tissue *t*, and *Z* denotes the z-score-transformation. The transformed expression values for each cell-type *(c)* are further summarized into cell type-specific marker-enrichment-score as the normalized sum of all the individual genes supporting a cell-type: 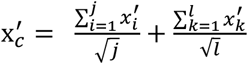, where *X*′ is the unique column of *X*′ corresponding to one cell, *c* is the specific cell-type within the tissue, and *i*,…,*j* are the indices corresponding to cell-type-specific marker genes, while *k*,…,*l* are the indices of negative marker genes that are not expected to be expressed in the cell type. Such transformation results in normalized expression matrix of c-by-n dimension, where each row represents one of the cell types and each column represents an individual cell. Finally, by summing up the values of each row (cell type) across the cells corresponding to a specific cluster *p*, the cluster summary enrichment-score (called ScType score) for each cell-type is calculated: 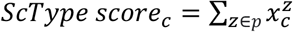. A cell-type with the highest ScType score is used for assignment to the cluster *p*. Such formulation guarantees marker gene specificity across both the cell-types 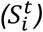 and cell clusters 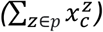 (Fig 1b-c), thus allowing for a highly accurate cell-type annotation. In addition to the cell-type assignments, the ScType web-portal (http://sctype.app) allows users to view the metadata based on which the assignment was made, view the markers that are enriched in each specific cluster, and plot the cumulative gene-specificity for different cell types as bar graphs.

### Publicly available datasets

In order to benchmark the ScType against the other cell-type annotation methods, we utilized six scRNA-seq datasets from public domain and re-analysed these using ScType platform. Five datasets were downloaded from Gene Express Omnibus (GEO): Human Liver (GSE124395), Human Brain (GSE67835), Human Pancreas (GSE85241), Mouse Lung (GSE63269) and Mouse Retina (GSE63473). Human PBMC dataset was downloaded from the 10x Genomics Dataset Repository (https://s3-us-west-2.amazonaws.com/10x.files/samples/cell/pbmc3k/pbmc3k_filtered_gene_bc_matrices.tar.gz).

### Comparison with other cell-type annotation tools

We compared the accuracy, running time and requirement of hardware resources of ScType against scSorter^30^, SCINA^33^, and scCATCH^34^ using the six scRNA-seq datasets (see Publicly available datasets). We used the default parameters to run all the methods. To provide unbiased comparisons, we used the same cell-type marker gene information (based on our ScType database) in scSorter and SCINA. scCATCH has its own in-built database that was used in the analysis.

### SNV identification using single-cell RNA sequencing

ScType utilizes raw scRNA-seq data to identify single-nucleotide variants (SNVs) present in each cell. ScType processes the raw input scRNA-seq BAM file using samtools^51^ and call the SNVs using Strelka2^52^. Next, ScType connects the SNV to each cluster using vartrix (https://github.com/10XGenomics/vartrix). As an extended feature, ScType also calculates the sum of total number of SNV present in the COSMIC cancer census genes^53^ as the combined SNV score (ScType_SNV score) for each cluster in a cancerous tissue profile. More specifically, ScType_SNV score summarizes the number of point mutations in the cancer genes for each cell type, calculated as the percentage of SNVs above the median SNV value within the particular cell cluster. ScType also incorporates recently implemented Bayesian segmentation approach, called CopyKAT^36^, to distinguish between aneuploid and diploid cells. Based on these two analysis, ScType automatically makes a classification between malignant and non-malignant cells. ScType assigns cells as non-malignant if ScType_SNV score < 20 and the majority of cell within the cell-type (>50%) are classified as diploid and the others as malignant. Users can use these two analyses to assign healthy and cancerous cell-type labels to the cell clusters, based on the assumption that cancerous cell clusters tend to have more SNVs in the cancer genes and aneuploidy is common for most human tumors^36^. The list of cancer consensus genes was downloaded from the catalogue of somatic mutation in cancer database (COSMIC v92)^53^.

### Code and data availability

The R source-code of the ScType algorithm is freely available at https://sctype.app/source_code.php to allow reproduction of the results and its further comparison against or integration with other algorithms. ScType is also freely available as an interactive web-tool at http://sctype.app. The ScType database is freely available at https://sctype.app/database.php.

## SUPPLEMENTARY FIGURES

**Supplementary Figure 1.**
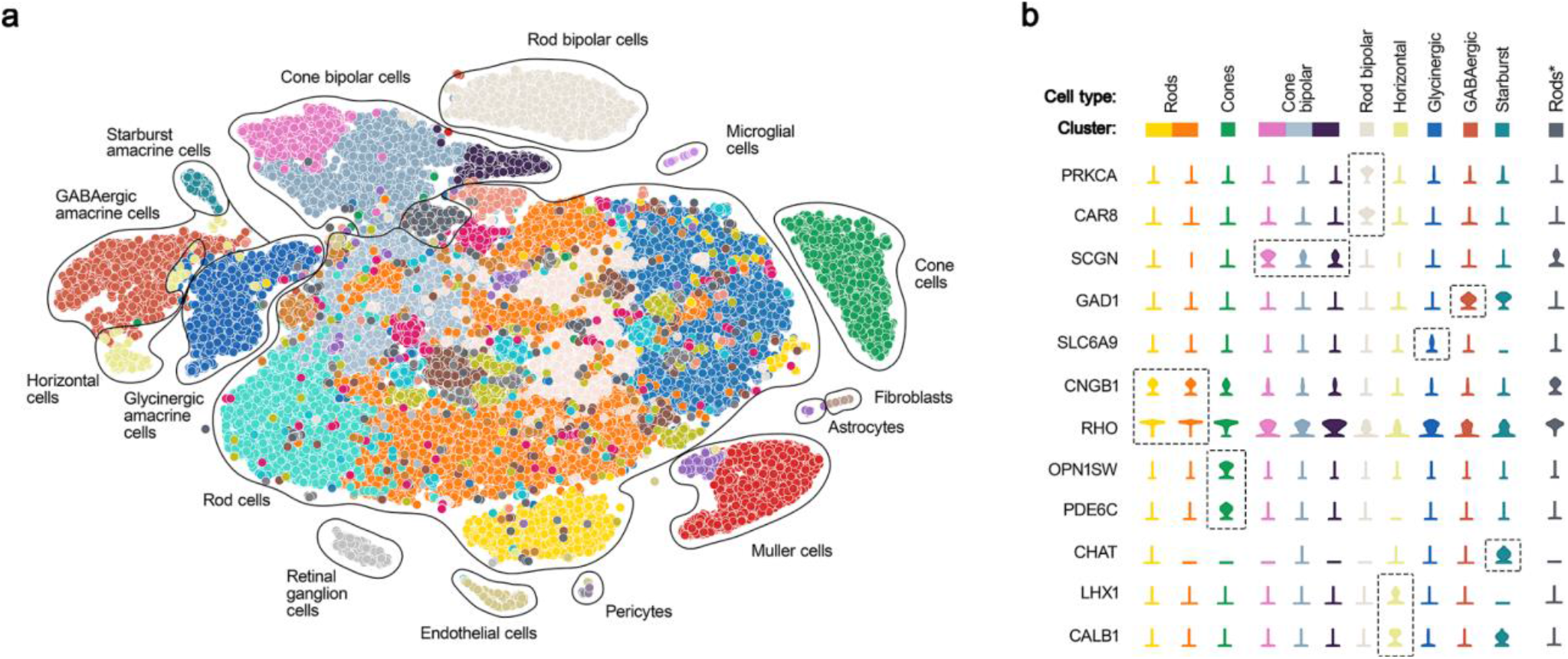
ScType cell-type annotation of mouse retina scRNAseq data^23^. **(a)** UMAP plot shows the automated cell-type annotations by ScType. **(b)** Violin plots show the expression levels of the high-specificity marker genes that were used as a validation of correct cell-type assignments.

**Supplementary Figure 2.**
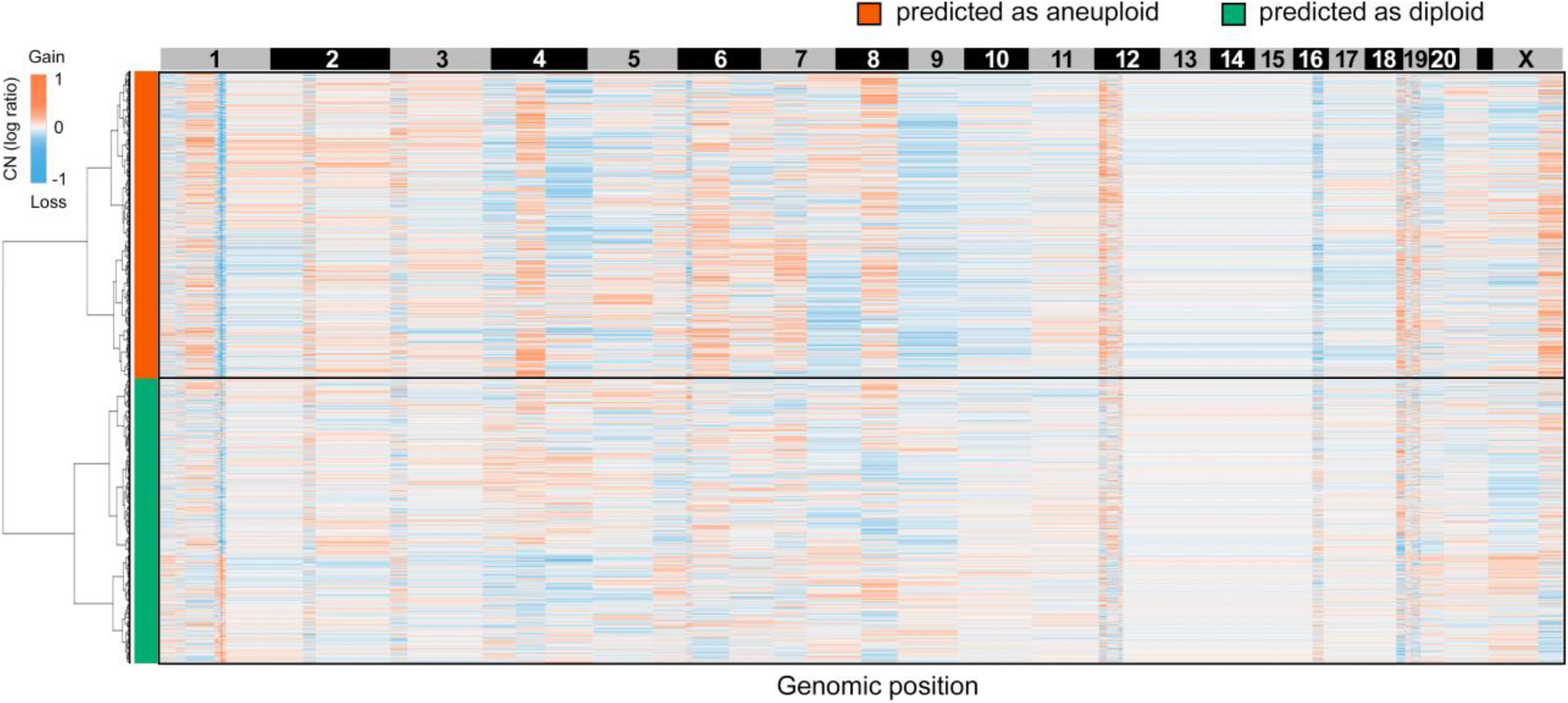
Clustered heatmap of single-cell copy number profiles estimated by CopyKAT^36^ tool in the AML patient sample.

## Notes

### Competing Interest Statement

The authors have declared no competing interest.

